# The interplay between stiffness and hyperglycemia on diabetic foot ulcer wound closure

**DOI:** 10.1101/2025.05.16.654387

**Authors:** Nourhan Albeltagy, Jennifer Patten, Karin Wang

## Abstract

**Introduction:** Diabetic foot ulcers are open wounds with impaired wound closure at the bottom of the foot. Although diabetic plantar skin is stiffer, which should enhance fibroblast mechanotransduction, fibroblasts still fail to migrate effectively. This suggests impaired wound closure is driven by another factor; hyperglycemia (≥11.1 mM glucose), which alters fibroblast mechanotransduction.

**Purpose:** To mimic diabetic foot ulcers by developing a 2D circular in vitro wound closure model system to investigate fibroblast mechanoresponse under diabetic plantar skin stiffness and hyperglycemia.

**Methods:** Polydimethylsiloxane was used as substrate, fabricated at 57 kPa and 90 kPa for normal and diabetic plantar skin stiffnesses, respectively. Cell culture media contained a 5.5 mM glucose concentration simulating normal blood glucose or an altered 11.1 mM glucose concentration simulating hyperglycemia.

**Results:** Time-lapse fluorescent imaging of wound assays reveals a restrictive effect of higher stiffness on migrating fibroblasts under normal glucose conditions, and a biphasic response to hyperglycemic conditions. Fibroblasts migrating on softer substrates mimicking normal plantar skin stiffness and under hyperglycemia have decreased velocity as predicted. Whereas cells migrating on stiffer substrates mimicking diabetic plantar skin stiffness and under hyperglycemia demonstrate increased cell velocity, overcoming the higher stiffness’s restrictive effect. Despite faster cell velocities on higher stiffness, wounds under normal glucose conditions still close faster than those under hyperglycemic conditions.

**Conclusion:** This research establishes a wound closure model demonstrating significantly slower wound closure in diabetic plantar skin with higher stiffness and hyperglycemic glucose levels compared to normal plantar skin with softer stiffness and normal glucose levels.

## Introduction

Diabetes, a dangerous metabolic disorder that impairs glucose regulation, is estimated to affect 700 million by 2045, up from 463 million in 2019. This places an unsustainable strain on slower-growing healthcare budgets [1]. A prominent risk of diabetes is the development of diabetic foot ulcers (DFUs), caused by diabetes-induced peripheral neuropathy and poor self-care. DFUs are chronic wounds that can lead to foot infections and amputations, which escalates the already costly $10,209 individual patient’s treatment to a pricey $78,069 per patient [2]. These added costs with DFUs complications account for a third of the global diabetes-related healthcare burden [3]. This indicates a need for more effective treatments driven by a deeper understanding of the mechanisms contributing to diabetic foot ulcers.

The development of DFUs is associated with physiological factors that hinder wound healing, such as a prominent inflammation response, the imbalance between extracellular matrix (ECM) degradation and assembly, and reduced fibroblast migration during wound healing in DFUs [4]. This research focuses on dermal fibroblasts as they are responsible for the contraction of the wound edges, remodeling the ECM, and closing wounds by migrating into wound sites [5–7]. The efficiency of wound closure is governed by fibroblast mechanotransduction [8, 9]. Mechanotransduction is the cellular mechanosensing and mechanoresponse that drives fibroblasts’ migration during wound closure [9, 10]. Some key mechanoresponses that can be evaluated and are affected by internal cellular mechanisms and regulations are: (i) cell velocity, which depends on actin polymerization and focal adhesion strength [11]; (ii) directionality, which is the ability of cells to maintain a persistent migration direction. It is influenced by lamellipodia extension at the cell’s leading edge [12]. And (iii) intrinsic actin alignment, which maintains directed migration and contraction forces [12].

Diabetic plantar skin exhibits higher stiffness compared to healthy skin [13]. Stiffness enhances fibroblast mechanotransduction by increasing focal adhesion formation and strength to drive actin contraction [14–16]. Yet, fibroblasts still fail to migrate effectively into diabetic foot ulcers. This impaired migration could be attributed to another main diabetic factor, hyperglycemia. Clinically, an individual is considered hyperglycemic if their random blood glucose levels are 11.1 mM or higher [17]. Higher glucose levels can alter various mechanotransduction pathways. Some proteins of interest can be the YAP/TAZ nuclear factors that regulate gene expression involved in cytoskeletal remodeling and focal adhesion dynamics in cell migration. And the Rac and Rho small GTPases that have a role in regulating the forward migration of cells [18]. Hyperglycemia reduces YAP transcription factor levels, inhibits alpha-smooth actin assembly, and decreases Rac1 activation [19–21]. These mechanisms were associated with deficient mechanotransduction, leading to impaired wound healing. High stiffness is also associated with higher Rac1 activation, YAP expression levels, and the number of actin stress fibers [22–24] However, the combined effect of hyperglycemia and higher diabetic plantar skin stiffness is unclear.

The most utilized technique for studying cell mechanoresponses *in vitro* is the scratch wound assay [25]. One type of scratch wound is the circular assay [26–28], which is more relevant to DFUs. Foot ulcers, unlike laceration wounds, tend to have rounded edges and are commonly circular [29]. In addition, wound assay curvature affects wound closure rate [30, 31]. Hence, a circular wound closure assay would more accurately model DFUs. Another factor to consider is the glucose level in the media. While research papers usually use high-glucose DMEM media 5 g/l (25 mM) to represent hyperglycemic conditions [32–36], the range for clinical cases starts at 11.1 mM. Therefore, 2 g/l (11.1 mM) glucose levels was used to model hyperglycemia, to more accurately represent clinically relevant glucose levels. Finally, plantar skin stiffness was determined by collecting data from literature averages that reported the plantar skin stiffness of live patients using the same ultrasonic method [37–40]. The stiffness was evaluated by testing the plantar skin areas susceptible to the highest-pressure forces during gait: the big toe, metatarsal heads, and heel [41].

Establishing an accurate model of DFUs with associated mechanoresponses will provide a better method of testing prospective therapeutics’ effects on DFU healing. Therefore, this study aims to establish a 2D circular *in vitro* wound closure model system simulating a diabetic foot ulcer to investigate the impact of diabetic plantar stiffness and hyperglycemia on the mechanotransduction of human dermal fibroblasts; specifically by analyzing cell velocity, directionality, and actin alignment during wound closure.

## Material and methods

### Cell line and culture media

Adult Human Dermal Fibroblasts (HDFa) acquired through The American Type Culture Collection (<u>PCS-201-012 | ATCC</u>) were maintained in Dulbecco’s Modified Eagle Medium (DMEM) with 10% FBS and 1% Penicillin-Streptomycin. Passages between 9-10 were used. Low-glucose DMEM 1g/L (5 mM Glucose) (CAT#11885084) was used to culture cells under normal glucose conditions (Ng). Separately, low-glucose DMEM media was supplemented with 2.5 ml of 200 g/L glucose solution (CAT#A2494001) to achieve the desired 2 g/L (11.1 mM Glucose) to culture cells under clinical diabetic glucose conditions (Dg). Each cell culture condition was passaged five times to adapt HDFas to the glucose levels and maintain a stable phenotype.

### Substrate fabrication and wound assay design

Table 1 shows plantar skin stiffnesses from the cited literature [37–40], demonstrated as Young’s Modulus, yielding an averaged stiffness of 56.0 ± 10.1 kPa for normal and 87.8 ± 5.9 kPa for diabetic plantar skin. Polydimethylsiloxane (PDMS; SYLGARD 184, CAT#50366794) was utilized to achieve the desired substrate stiffness by casting a thin layer (0.2 g) of PDMS in each well of a 12-well plate. Stiffness was evaluated via Bose ElectroForce 3200 compression test with 10 N compression force of a 50 N load cell, from the 10% displacement of ∼0.5 cm long x 2 cm wide PDMS cylindrical sections. a range of base-to-crosslinker ratios, (40:1) to (46:1), were tested for PDMS’s fabrication to decide on our target base-to-crosslinker ratio. After mixing, the PDMS mix was degassed for 1 hour, incubated at 60°C for 4 hours, then sterilized under UV for 15 minutes. 3 trials were tested per ratio.

**Table 1.** Stiffnesses of each section of plantar skin, adapted from the literature.

| Normal Plantar Young’s Modulus (kPa) |  |  |  |  |  | Diabetic Plantar Young’s Modulus (kPa) |  |  |  |  |  |  |
| --- | --- | --- | --- | --- | --- | --- | --- | --- | --- | --- | --- | --- |
| big toe | 1st MTH | 2nd MTH | 3rd MTH | 5th MTH | heel | big toe | 1st MTH | 2nd MTH | 3rd MTH | 5th MTH | heel | Ref. |
| 62.3 | 80.1 | - | 99.4 | 81.8 | 69.8 | 77.1 | 91.4 | - | 121.5 | 99.5 | 85.4 | [37] |
| 30.4 | 49.0 | - | 48.0 | 36.0 | 49.5 | 66.4 | 72.0 | - | 98.4 | 88.0 | 86.0 | [38] |
| 41.0 | 41.0 | 43.0 | - | - | 41.0 | 100.0 | 118.0 | 68.0 | - | - | 43.0 | [39] |
| 50.5 | 62.1 | 52.9 | - | - | 45.0 | 85.3 | 96.3 | 84.8 | - | - | 65.6 | [40] |
| Average = 56.0 ± 10.1 kPa |  |  |  |  |  | Average = 87.8 ± 5.9 kPa |  |  |  |  |  |  |

### Assay start and time-lapse imaging

**The model workflow and design are shown in Fig. 1. The PDMS substrates were coated with 30 µg/ml fibronectin to facilitate cell adhesion to the hydrophobic PDMS. 1×1 mm cylindrical PDMS masks were placed on the coated PDMS surface, followed by cell seeding at a 40k cells/cm^2^ density. Steps per 1 well are shown in**

**Fig. 1.**
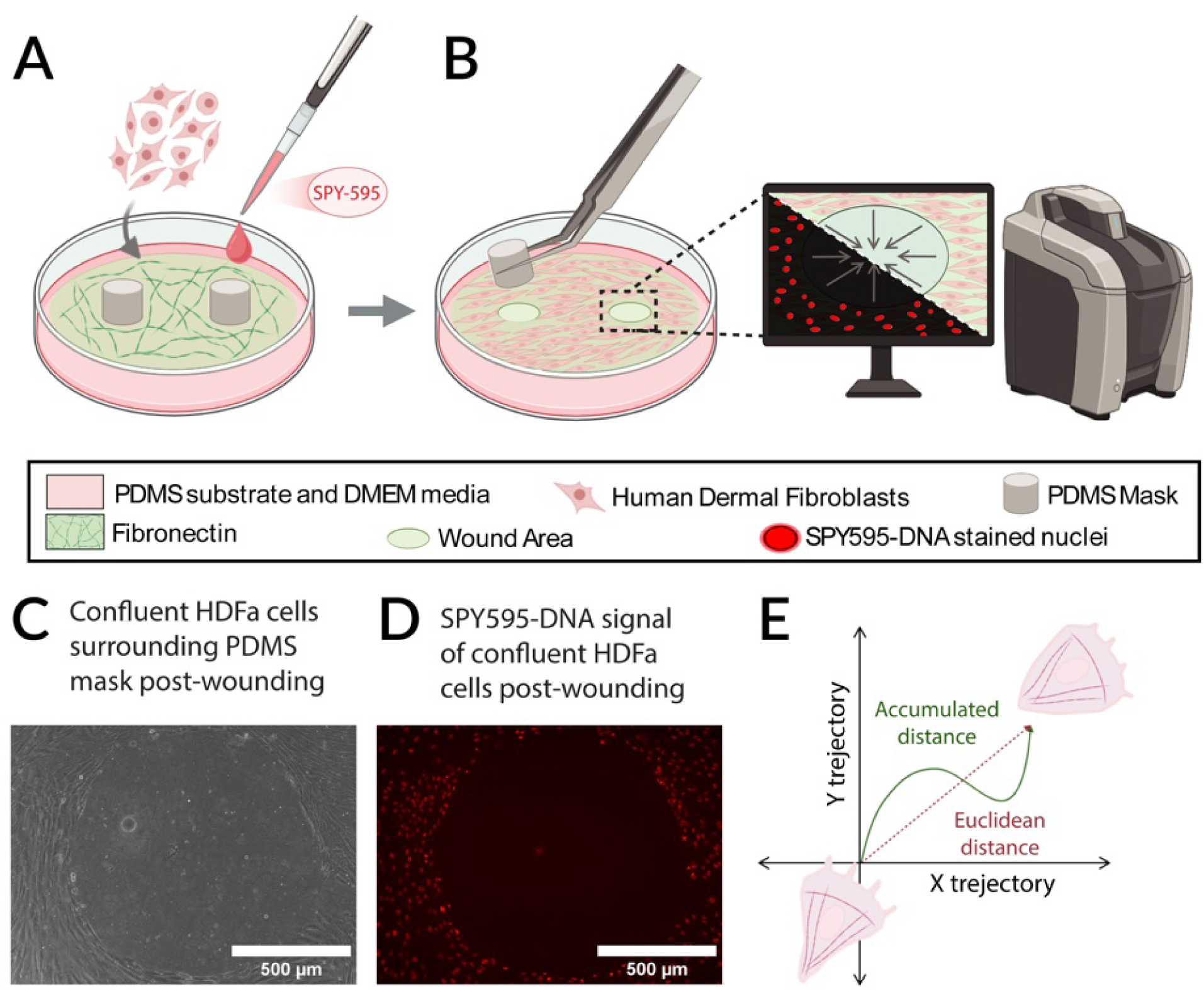
2D circular wound assay model system. A) PDMS substrates were coated with fibronectin before cell seeding, with 1 mm wide PDMS masks in prospective wound areas. HDFas’ nuclei were tagged with SPY595-DNA live stain before time-lapse microscopy on the Keyence. B) 18 hours after seeding, simulation of wounding was initiated by mask removal. Circular wounds were imaged for 48 hours at 10 min intervals, with phase contrast and TxRed channel imaging. C) Phase image of confluent cell layer around wound area created from cylindrical PDMS mask removal (post-wounding). D) SPY595-DNA stained nuclei signal of confluent cells around the wound area (post-wounding). E) The plot shows an example of a cell track in the XY plane, data acquired from the graph is used to calculate velocity and directionality; velocity = Euclidean distance/migration time, directionality = Accumulated distance/Euclidean distance. Accumulated distance is the total distance a cell travels, while euclidean distance is the distance between where the cell starts to where it ends. (A-B,E) Made with BioRender.

Fig. 1A, B. The cells were cultured overnight with SPY595-DNA live stain (CAT#CY-SC301) at a 1:3000 dilution before assay start.

**18 hours after seeding, cells reached confluency around the circular PDMS masks. After confluency confirmation, masks were removed to simulate wounding (**

**Fig. 1B). The plates were equilibrated in the imaging incubation chamber for 2 hours before time 0 to stabilize PDMS expansion or shrinkage in the new environment to achieve a stable focus on the Keyence BZ-X800 fluorescence microscope (RRID: SCR_023617). Time-lapse imaging was acquired with a 10x objective for 48 hours at 10-minute intervals. Cells were imaged in phase contrast and with a BZ-X Filter TexasRed filter to detect SPY595-stained cell nuclei. An example of the wound generated is shown in** Fig. 1C, D.

### Immunohistochemistry and actin alignment detection

After wound closure, HDFas were fixed with paraformaldehyde (PFA, CAT#AAJ61899AK, FisherScientific) and immunostained with DAPI (CAT#D1306), and Alexa Fluor 568 Phalloidin (CAT#A12380). Actin angle alignment data were detected using the Directionality function in ImageJ from the immunostained images. This function computes a histogram of orientations and fits a Gaussian curve to the highest peak. The F-actin filaments were color-coded depending on angle alignment with the ImageJ plugin, OrientationJ. Data was normalized with the max peak angle to 0 degrees and evaluated using the goodness of fit. The closer the goodness of fit is to 1, the more the data matches the normal distribution. A representative cell F-actin alignment is shown in Fig. 4B for reference.

### Wound closure assay

Wound closure rate was evaluated from the phase video by defining wound edges in ImageJ (RRID: SCR_003070) at 8-hour intervals. Each time point area was calculated by the following equation to plot the wound closure rate.

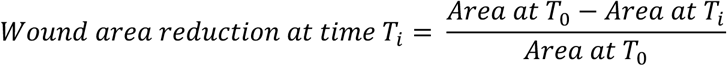

### Migration tracks detection and analysis

SPY595-DNA stained nuclei fluorescent time-lapse microscopy images were used to detect cell migration tracks via the ImageJ Trackmate plugin. The data acquired from Trackmate was plugged into Chemotaxis and Migration Tool 2.0 (http://www.ibidi.de/-applications/ap_chemo.html) to calculate individual cell velocity, directionality, and migration degree angle. The equations for the analysis are as follows:

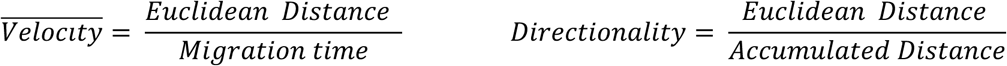

**Euclidean Distance is the shortest straight-line distance from where the cell started to where it ended in the XY plane, while the Accumulated distance is the total distance a cell migrated. An example is shown in**

Fig. 1E. For rose diagrams, a MATLAB function (supplementary code1) was modified from (Wind Rose,MATLAB) to plot cell data [42]. The rose diagram represents the distribution of cell migration direction and velocity. The migration angles were grouped into 15 bins each covering a 24-degree range. The height of each bin indicates the percentage of cells migrating in that direction while the color segments within each bin represent different velocity ranges. The trajectory of the migrated tracks and the center of mass (COM) displacement were plotted using a custom-made MATLAB code (supplementary code2). COM represents the center point of all tracks’ final coordinates, and the displacement of each center from the initial (0,0) coordinate is calculated via the following equation.

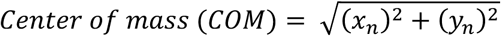

x_n_: final average x coordinate of all tracks, y_n_: final average y coordinate of all tracks.

### Statistical analysis

All statistical analyses were performed using GraphPad Prism 9. For wound closure, data was optimized for a 100% closure rate, where 0% closure at time zero, and 100% is wound closure. A line best-fit analysis was used to generate slope values. Two-way ANOVA with Tukey’s multiple comparison test was performed for velocity, directionality, and actin goodness of fit analysis, and a T-test was performed for stiffness differences. α = 0.05 for a 95.00% CI.

## Results

### Fabricated PDMS stiffnesses match target plantar skin stiffnesses

Table 2 shows the summary of the compression testing data of our 2 final ratios that best match literature averaged plantar skin stiffnesses in Table 1. Each fabricated PDMS sample was evaluated through compression testing. The data shows that PDMS crosslinking ratios: 1) 45:1 with Young’s modulus of 57.5±7.0 kPa simulates normal plantar skin stiffness (Ns), 2) 42:1 with Young’s modulus of 90.2±1.7 kPa, simulates diabetic plantar skin stiffness (Ds). Fig. 2A and Fig. 2B shows those acquired PDMS stiffnesses compared to the stiffness levels from the literature, and how they match the range and statistical differences of plantar skin stiffnesses in healthy and diabetic individuals shown in Table 1.

**Fig. 2.**
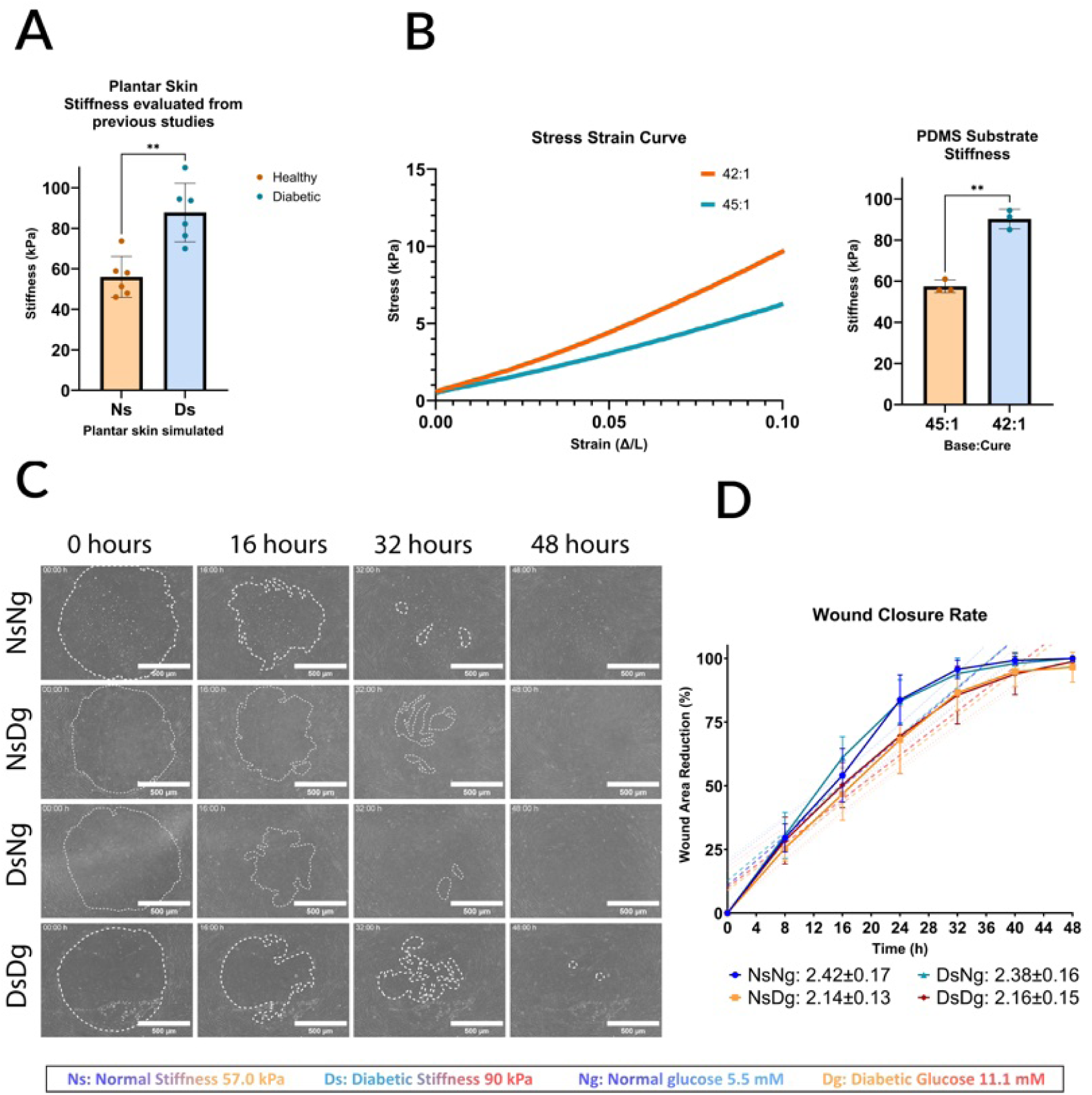
Our fabricated PDMS stiffness compared to the literature data. A) Data from Table 2 is shown in the graph. (Ns) Normal stiffness represents healthy patients’ plantar skin stiffness averages, (Ds) Diabetic stiffness represents diabetic patients’ plantar skin stiffness averages. B) Stress-strain curve of the fabricated PDMS substrates, evaluated via compression testing of 10% sample strain under 10 N load using Bose ElectroForce 3200. 42:1 base to crosslinker ratio had a Young’s modulus of 90±6 kPa to simulate diabetic plantar skin stiffness (Ds), and 45:1 had a Young’s modulus of 57±5 kPa to simulate normal plantar skin stiffness (Ns). Both (A) C (B) show significant differences between the 2 stiffness values,**P≤0.0020. C) One representative wound per condition is shown. Phase images show the wound area reduction at time points 0, 16, 32, 48 hours for our four conditions: Normal stiffness Normal glucose (NsNg), Normal stiffness diabetic glucose (NsDg), Diabetic stiffness normal glucose (DsNg), and Diabetic stiffness diabetic glucose (DsDg). Scale bar = 500 µm. D) The wound closure rate with best-fit line and standard error, made with GraphPad Prism, slopes are NsNg = 2.42±0.17 (R2=0.93), NsDg = 2.14±0.13 (R2=0.91), DsNg = 2.38±0.16 (R2=0.88), and DsDg = 2.16±0.15 (R2=0.98). N = 4.

**Table 2.** Compression testing results for the target Base-to-crosslinker ratio, values in kPa.

| (base: crosslinker) | (42:1) | (45:1) |
| --- | --- | --- |
| Trial 1 | $94.4 \pm 5.3$ | $55.8 \pm 5.0$ |
| Trial 2 | $91.3 \pm 16.4$ | $55.6 \pm 5.6$ |
| Trial 3 | $85.1 \pm 5.5$ | $61.1 \pm 10.8$ |
| Average (Mean ± SD (kPa)) | $90.2 \pm 10.7$ | $57.5 \pm 7.0$ |

### Wound closure rate is slower under the effect of hyperglycemia

Wound closure rates (Fig. 2D, Supplemental Fig. 1A, B, C and D) had slopes with standard error (SE) of: NsNg = 2.42±0.17 (R^2^=0.93), NsDg = 2.14±0.13 (R^2^=0.91), DsNg = 2.38±0.16 (R^2^=0.88), and DsDg = 2.16±0.15 (R^2^=0.98). Based on slope values, there is no significant difference in wound closure on normal and diabetic PDMS stiffnesses within normal glucose level conditions, as their values fall within each other’s SE. The same applies to wound closure on normal and diabetic PDMS stiffnesses within diabetic glucose level conditions. However, when comparing the wound closure slopes between normal and diabetic glucose level conditions on either stiffness, the values do not overlap. This suggests that the differences between the two glucose levels might play a significant role in altering wound closure.

### HDFas have a biphasic response to glucose levels under different stiffnesses

**Representative wounds from each condition are shown in**

**Fig. 3. For all conditions, HDFas migrated towards the center of the wound at varying velocities. HDFas detected closer to the wound edge migrate faster, while cells that begin migrating later into the wound area moved at a slower speed. This observation aligns with leader-follower dynamics seen in cells’ collective migration** [43, 44]**. This distribution can be seen further in the rose diagrams (**

**Fig. 3.**
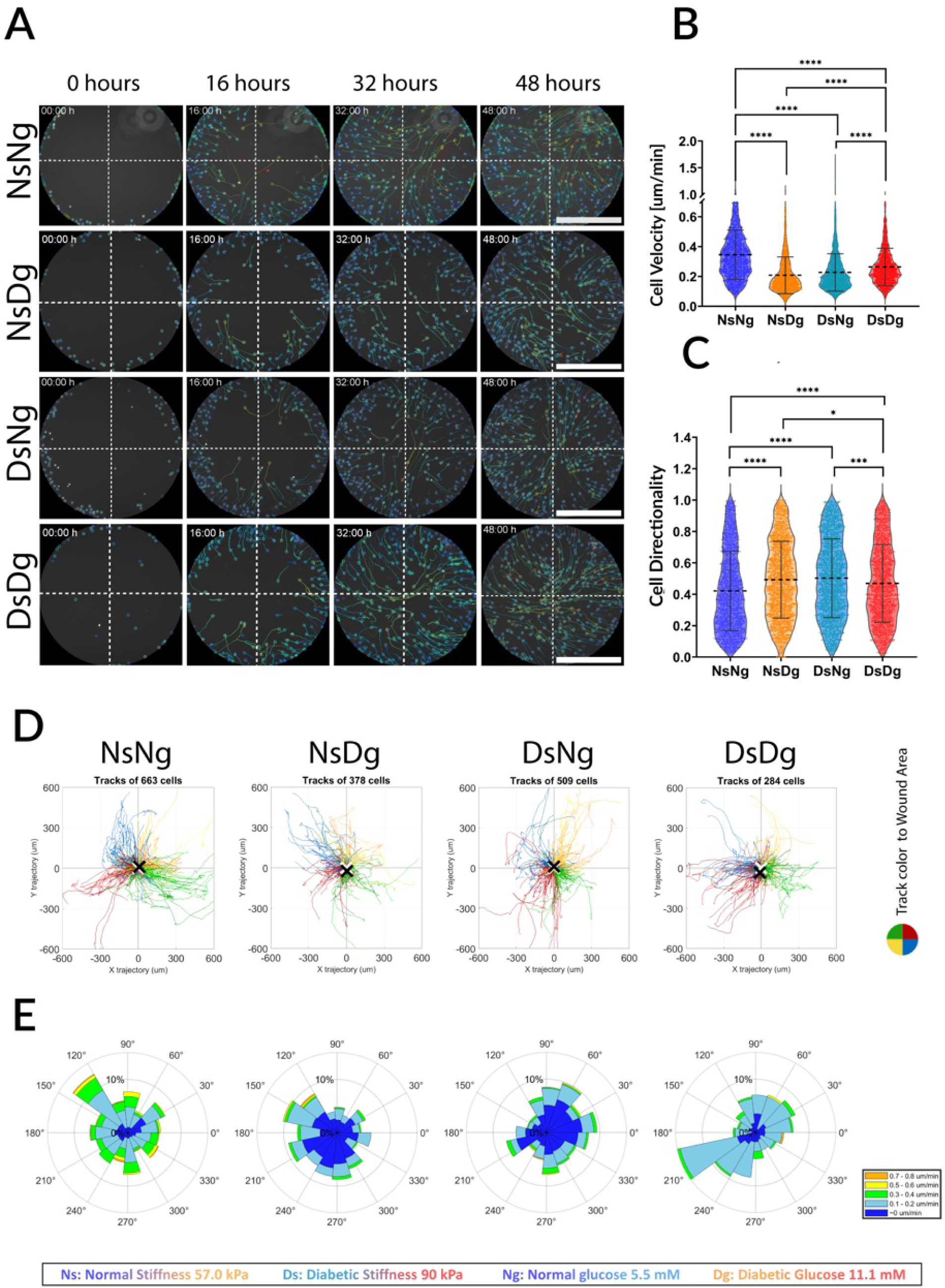
Migration dynamics and cell distribution. All figures are from one representative wound per condition. A) TrackMate videos of cell migration tracks at time points (0, 16, 32, 48) hours. The tracks are color-coded according to track mean speed, as shown in the speed range [μm/sec]. Scale bar = 500 μm. B) Cell migration dynamics, evaluated via GraphPad Prism, show: i) NsNg velocity was significantly higher than NsDg, DsNg, and DsDg. ii) DsDg velocity was significantly higher than NsDg and DsNg. This shows how stiffness decreased velocity in normal glucose but increased it in diabetic glucose, while diabetic glucose decreased velocity in normal stiffness but increased it in diabetic stiffness. Results represent a biphasic cellular response to the combination of stiffness and glucose level factors. C) for directionality, i) NsNg directionality was significantly lower than all other conditions, and NsDg was significantly higher than DsDg, ii) DsDg was significantly lower than DsNg and NsDg. This reveals an inverse relationship between the velocity and directionality of cell migration. *P≤0.0500, **P≤0.0050, ***P≤0.0005, **** P<0.0001. Number of cells per condition: NsNg = 1859, NsDg = 2169, DsNg = 2217 and DsDg = 1673. D) Graphs generated through MATLAB show the homogeneity of cell migration trajectories after 48 hours. The associated number of tracks detected is also indicated. The color code represents which quarter of the wound area the cell started from: red is the upper right corner, green is the upper left corner, blue is the lower right corner, and yellow is the lower left corner. A black X shows the center of mass (COM) deviated location for each wound, NsNg COM = 13.61 μm, NsDg COM = 23.84 μm, DsNg COM = 10.93 μm, and DsDg COM = 35.12 μm. All COMs were less than 10% deviation from the wound radius of 500 μm. E) The rose diagrams show the distribution of migration direction represented in angles. Each data point was divided into 15 bins, and each bin was divided into the percentage of cell velocities that moved into the direction of the respective bin. Data showed a small fraction of faster moving cells per direction, followed by a bigger fraction of slower moving cells.

Fig. 3E) Each bin in the rose diagram shows cell speeds across three or more ranges, with the highest speeds (>0.3 μm/min and >0.5 μm/min) occupying small fractions of total cells per bin. This pattern suggests a small number of cells initially migrate rapidly into the wound area, followed by a larger number of slower cells.

**Trackmate videos are shown in Supplemental Fig. 2A, B, C, and D, and time points captures are shown in Fig. 3A. The corresponding Trajectory graphs of all conditions (**

Fig. 3**Error! Reference source not found.**D) demonstrate how all migrating cells moved towards the wound center in all conditions. The center of mass (COM) was detected as the average of all final migration points. When the center of mass (COM) of migrating cells falls within the central region of the wound, it indicates uniform and coordinated migration from all wound edges toward the center. To evaluate this, the wound area can be divided into concentric circular zones based on fractions of the original wound radius. For example, in a wound with a 500 μm radius, a COM within 10% of the radius (i.e., within 50 μm of the center) suggests that cells at the wound edge began migrating simultaneously and evenly at time 0. Conversely, a COM beyond 50% of the radius (i.e., farther than 250 μm from the center) would indicate that migration either did not start uniformly across all wound edges or was not directed towards the wound center. In this study, all detected COM values remained within the 10% threshold, indicating consistently centralized migration initiation. The data showed COM of: NsNg = 13.61 μm, NsDg = 23.84 μm, DsNg = 10.93 μm, and DsDg = 35.12 μm. This reflects how cells collectively migrated to fill and close the wound area in all four of our conditions, despite any hindering effect of diabetic glucose levels.

Individual HDFa migration dynamics were evaluated from at least 3 wounds for each condition (Fig. 3B). Diabetic glucose level significantly decreased the velocity of migrating cells on normal stiffness (57 kPa). However, on diabetic stiffness (90 kPa), the same diabetic glucose level increased the velocity of migrating cells more than that of cells exposed to normal glucose levels. The higher diabetic stiffness decreased cell velocity under normal glucose levels (5.5 mM) more than diabetic glucose conditions (11.1 mM). Where cell velocity on diabetic stiffness and under normal glucose conditions (DsNg) was significantly lower than that of cells on diabetic stiffness and under diabetic glucose conditions (DsDg). This showcases a biphasic response of HDFa to hyperglycemia, depending on the underlying substrate stiffness.

Cells on normal stiffness and exposed to normal glucose conditions (NsNg), which migrated the fastest, exhibited the lowest directionality or the most random migration among all conditions. This is not the first instance where velocity and directionality in collective migration were found to be non-proportional [45, 46]. On higher diabetic stiffness, this relationship between wound closure rate, velocity, and directionality was reversed. Cells migrating on higher stiffness and exposed to normal glucose levels (DsNg) had higher directionality than cells exposed to diabetic glucose levels (DsDg). And despite the cells’ lower velocity on diabetic stiffness while exposed to normal glucose levels (DsNg), they had faster wound closure rates.

### F-actin alignment under the combined effect of diabetic glucose and diabetic stiffness

The alignment of F-actin showed no significance in the goodness of fit between all the different conditions (Fig. 4C). This was also shown by how actin alignment showed almost identical normal distribution curves (Fig. 4D). This means that cells can still maintain their internal polarization and normal rate of actin polymerization at the leading edge in the hyperglycemic conditions represented as diabetic glucose level.

**Fig. 4.**
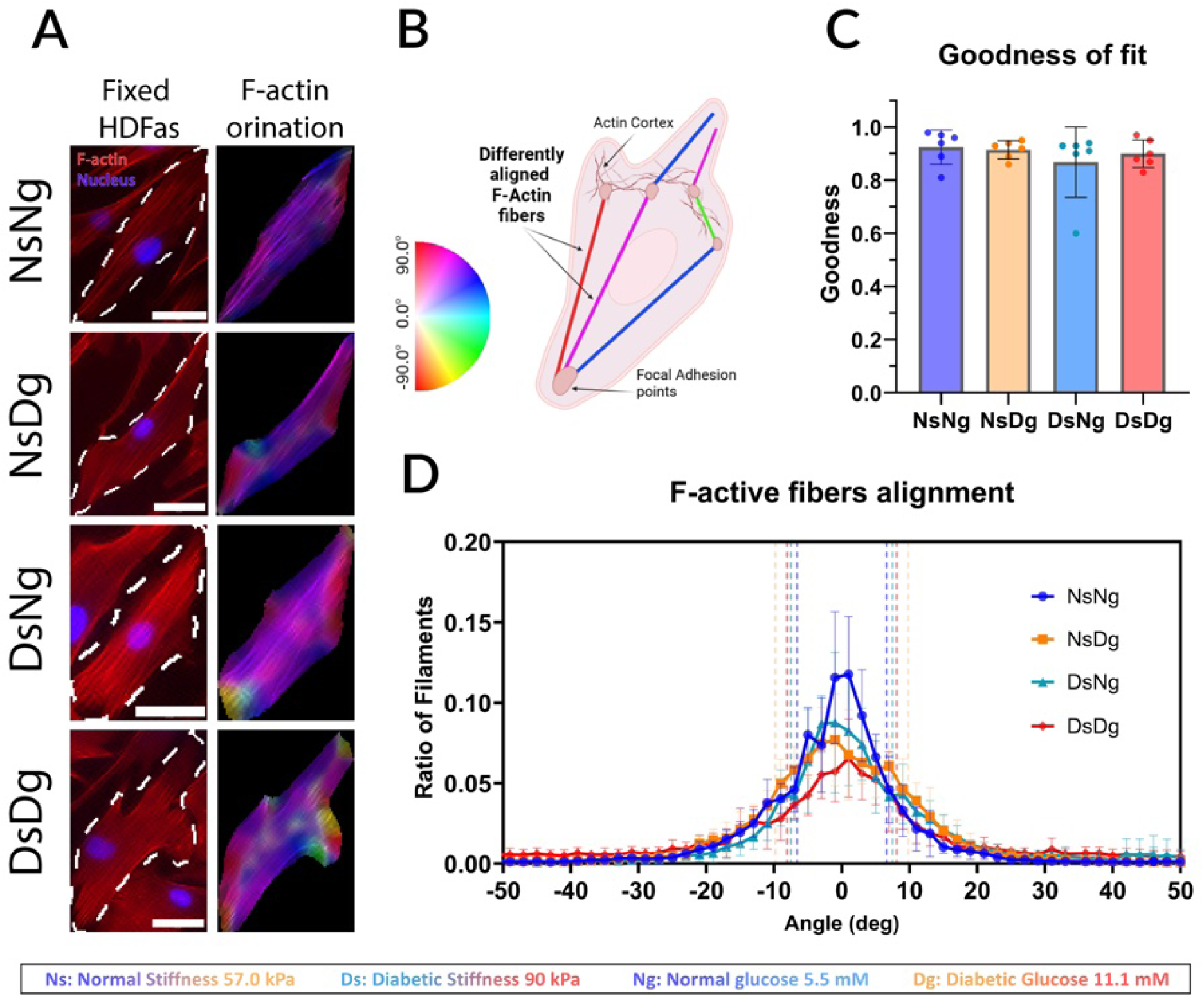
Actin filaments orientation evaluation from immunohistochemistry, A) One HDFA cell example with is shown per condition. Post 48-hour migration, cells were stained with phalloidin (red) to detect actin and DAPI (blue) to detect the cell nuclei. Each cell have the respective orientation map that is color-coded depending on the color-wheel guide to show the orientation angle of each filament. Filaments following the same orientation would show a more homogenous color range. Scale Bar = 50 um B) An illustration showing the F-actin fibers extending between focal adhesion points within a migrating cell, each fiber is color coded according to the guide for elaboration on the F-actin orientation map in (A). C) Fiber alignment curve goodness of fit. D)The graph shows the percentage of total filaments per image crop following each orientation angle, normalized to 0 degrees. With SD of each curve represented in a dotted line.

## Discussion

Our fabricated PDMS stiffnesses were optimal substrates to model normal and diabetic plantar skin stiffnesses (Fig. 2A, B) similar to plantar skin stiffness from clinical studies [37–40]. The achieved stiffness of 58±7 kPa from the ratio 45:1 (Base: crosslinker) was the closest to the target 56±10 kPa from healthy patients’ plantar skin stiffness. And the stiffness of 90±11 kPa achieved with 42:1 was the closest to the target 88±6 kPa from diabetic patients’ plantar skin stiffness. This ensures the accuracy of the model to the clinical cases of DFU.

After performing the wound assay and measuring wound closure, we observed a significant difference in wound closure between healthy conditions with normal glucose levels of 5.5 mM and hyperglycemic levels of 11.1 mM. But no difference in wound closure rate was observed between the two stiffness values mimicking normal and diabetic plantar skin stiffnesses. Previously, hyperglycemic 11.1mM glucose concentrations have been shown to decrease wound closure rate; as a previous study by Pahwa et al demonstrated that a glucose concentration of 11.1 mM was enough to slow mice osteoblasts cell migration in the scratch wound assay [47]. This confirms how hyperglycemic blood glucose levels alone have an adverse effect on different cells’ migration abilities. These results also further confirm that stiffness alone does not have a significant effect on wound closure rate [48]. However, stiffness does affect cell mechanoresponses not only glucose concentrations; therefore, we further investigated individual cell dynamics and actin alignment to unravel the underlying cellular responses.

Collective cell migration encompasses the coordination between migrating cells while maintaining cell-to-cell contact, and those migrating cells have differing roles, such as leader and follower [49, 50]. Leader cells that migrate first and closer to the wound edges are faster and followed by slower moving follower cells [43, 44]. In high cell densities of migrating fibroblasts, cells collide during migration, forming transient cell contacts and overlapping lamellipodia. This collision causes a directional change in migrating cells [51–54]. Park et al termed this behavior as “cell collision guidance” [53]. When fibroblasts are exhibiting this transient collective migration behavior, the first line of cells takes the leader role as they extend their leading edge forward, maintaining a connection to cells at the rear [54]. Those dynamics were represented in our circular wound closure model, observed in

Fig. 3D, E. More cells follow the path of faster cells, leading to an unequal distribution of migration across the area. Cell collision guidance from high densities maintained a general migration direction towards the wound center, as shown by the low COM values. Our data verified this cell behavior in

Fig. 3. This demonstrates that despite hyperglycemic conditions, cells were able to maintain their usual collective migration behavior, but what differed was cell velocities. The rose diagrams of cells on normal stiffness exposed to normal glucose levels (NsNg) and on diabetic stiffness exposed to diabetic glucose levels (DsDg) exhibited a higher fraction of faster cells per bin than the more equally distributed cells on normal stiffness exposed to diabetic glucose levels (NsDg) and on diabetic stiffness exposed to normal glucose levels (DsNg). This can be related to how HDFas migrated faster on normal stiffness with normal glucose (NsNg) than with diabetic glucose (NsDg), but migrated faster on higher diabetic stiffness with diabetic glucose (DsDg) than with normal glucose (DsNg), showing an opposite response to hyperglycemia under the two stiffnesses.

While previous literature focused on wound closure rate in scratch assays as the main indicator to test various cellular mechanisms [47, 55–57], this study investigates individual cell dynamics as an approach to identify key mechanoresponses contributing to the deficient wound healing response in DFU. The first observation was an inverted relationship between cell velocity and directionality. However, slower cell velocity does not necessarily translate to slower wound closure in all cases presented here. A persistent migration towards the wound center achieves faster coverage of the wound area, even if cells are migrating more slowly. This has been observed before [45, 46], and can be further explained by the increased number and size of focal points, and actin polymerization under the effect of higher stiffness.

In this model system, HDFa cells migrated faster and more randomly under hyperglycemic conditions, but only when coupled with the higher diabetic stiffness. Looking at higher diabetic stiffness alone, we observed that, unlike the predicted positive effect of higher stiffness on cell migration, cell velocity significantly decreased while migrating on higher diabetic stiffness. The decreased velocity resulting from increasing substrate stiffness has been observed previously in fibroblasts [58] and various other cell types [59–64], and some other studies reported no velocity change [65, 66]. One consistent finding between those studies was the increase in persistence and directionality with higher stiffness [58, 59, 65, 66] which matches our findings. In Yu. et al [64], mast cells’ optimal cell velocity was observed on 51.25 kPa substrate stiffness, and this velocity decreased on substrate stiffnesses of 83.28 kPa. This range is comparable to the substrate stiffnesses in our study (57 kPa and 90 kPa) and confirms our findings that higher stiffness beyond an optimal range decreased HDFa migration velocity. This is likely due to the fact that increased substrate stiffness leads to stronger and larger adhesions. Those enhanced adhesions allow cells to better adhere to the substrate, helping them to stay on course, but also cause restricted migration due to rigid connections in the trailing edge [15, 16, 67]. Ji et al called this the ‘motility factor,’ which is the ratio of the driving force to cell movement resistance. This motility factor also drives the inverse relationship between velocity and directionality, as enhanced adhesions also induce a more aligned internal cell polarity [16].

Further looking into the effect of hyperglycemia, although previous literature reported a decrease in velocity at higher glucose levels (∼25 mM) [20, 47, 68, 69], HDFa under 11.1 mM showed a biphasic response to hyperglycemia, dependent on substrate stiffness. Other types of cells, such as endothelial cells, showed a similar biphasic response, where cells had a higher velocity under high glucose when hypoxic concentrations were absent [70]. And biphasic responses in HDFa have been observed previously for different stimuli [71–74]. In our results, hyperglycemia increased cell velocity under the restrictive effect of the higher diabetic stiffness (DsDg), while on normal stiffness (NsDg), it decreased velocity as predicted. A possible explanation is that glucose availability allowed cells to produce more energy and force through glycolysis, providing ATP for the polymerization and depolymerization of F-actin [75–77]. The provided energy from the excess of glucose in this model supported cytoskeletal rearrangements to overcome the higher stiffness’s restrictive effect. However, this increase in velocity and decrease in directionality in cells under hyperglycemia with diabetic stiffness (DsDg) did not match that of cells exposed to normal glucose and normal stiffness (NsNg), which may be why those faster migrating cells in hyperglycemia still did not similarly close wounds as rapidly. Wounds with higher diabetic stiffness and under hyperglycemic conditions still have a slower wound closure rate than wounds with normal stiffness and under normal glucose conditions.

There was no significant difference in the goodness of fit between all the actin alignment different conditions. In a previous paper by Xing et al. [21], F-actin alignment was shown to be significantly decreased under a higher glucose level of 25 mM, but the same effect was not observed in our migrating cells exposed to 11.1 mM glucose levels. This likely indicates that cells can still maintain their internal polarization of actin fibers despite the hyperglycemic glucose level of only 11.1 mM. This leads us to suspect that more intrinsic factors are at play, causing the decreased wound closure rate under hyperglycemia. Even if cellular mechanoresponses were completely reversed, that will not always reflect the same in wound closure rate. Different combinations of mechanoresponses can lead to the same final effect on wound healing, but it would indicate a different mechanotransduction pathway activation and deactivation.

This presented model has the potential to be a suitable testing environment for investigating the mechanotransduction pathways altered in DFU. A future candidate for investigating mechanotransduction pathways could be the Rac1 protein, as it mediates lamellipodia extension and cell migration directionality at the leading edge of migrating cells [78]. Rac1 is also a downstream effector of both high glucose and high stiffness [21, 23, 79]. and is one cause of decreased wound healing in diabetes [21].

It is also important to mention that this model closely simulates the diabetic foot ulcer environment in a 2D context. Although a 3D model incorporating wound depth and extracellular matrix components would provide additional spatial cues and binding sites for fibroblasts, adding more factors also increases the complexity of isolating cellular responses to individual stimuli.

## Conclusion

This study highlights the importance of evaluating mechanosensing and migration responses at the cellular level rather than relying solely on gross wound closure measurements. The biphasic response of fibroblast migration, as demonstrated here, can only be detected through cell-specific evaluation of velocity and directionality. These findings suggest that fibroblast migration in diabetic wounds is influenced by complex biochemical and mechanical interactions, reinforcing the necessity of a multifactorial approach in wound healing research. Notably, while cell velocity and directionality patterns were reversed on higher diabetic stiffness, lower glucose conditions were consistent in promoting faster wound closure than hyperglycemic conditions. This suggests that the mechanotransduction pathways activated or deactivated vary by substrate stiffness, emphasizing the critical role of substrate stiffness in cellular migration studies.

## Acknowledgement

The authors acknowledge the shared equipment used in the Bioengineering department at Temple University. We thank BioRender for providing a platform to create parts of the schematics used in the figures.

## Funds

N.A was supported by the Fulbright Master’s Scholarship program and Teaching Assistantship from the Bioengineering Department at Temple University. K.W. acknowledges startup funds from the Bioengineering Department at Temple University, an Office of the Vice President for Research Bridge grant, and a grant (#2305) from the W.W. Smith Charitable Trust

## Statements and Declarations

### Contributions

NA: Conceptualization, Methodology, Validation, Data Analysis, Visualization, Writing. JP: Methodology, Validation, Review and Editing. KW: Conceptualization, Methodology, Validation, Visualization, Writing, Funding acquisition, Supervision.

### Conflict of interest

The author declares no conflict of interest.

### Human and Animal Rights

This study did not involve human subjects or animal research.

## Notes

### Competing Interest Statement

The authors have declared no competing interest.

