## Supplementary Information for "The interplay between stiffness and hyperglycemia on diabetic foot ulcer wound closure"

### Supplementary codes:

Code1: Rose is adapted from (Wind Rose, MATLAB). Cellrose function Generates a polar histogram (rose plot) of cell movement directions and speeds. Inputs include a vector of cell's migration angle, and a vector of cell velocities. Output is a 2D histogram matrix representing the percentage of cells in each direction and speed bin. Supplementary include cellrose.m function and cellrose_script.m example to run the function.

Code2: Migration tracks script. This MATLAB code imports cell migration data, normalizes and plots individual cell trajectories on a 2D graph, color-coding each track based on its starting quadrant. It also calculates and marks the center of mass of all cell endpoints, providing a visual summary of cell movement patterns over 24 hours. Input is an .xlsx file in the format defined by (ibidi Chemotaxis and Migration Tool, [Chemotaxis and Migration Tool | Free Software | ibidi])

### Supplementary figures:

**Supplemental Fig. 1 Representative wound closure videos.** Wound area reduction was evaluated at time points 0, 8, 16, 24, 32, 40, and 48 hours. Phase images were acquired with a Keyence BZ-X800 fluorescence microscope and a 10x objective. The two substrate stiffnesses are normal stiffness (Ns) at 57±5 kPa and diabetic stiffness (Ds) at 90±6 kPa. The two glucose conditions are normal glucose levels (Ng) at 5.5 mM, and diabetic glucose levels (Dg) at 11.1 mM. A) Wound closure on normal stiffness with normal glucose levels (NsNg). B) Wound closure on normal stiffness with diabetic glucose levels (NsDg). C) Wound closure on diabetic stiffness with normal glucose levels (DsNg). B) Wound closure on diabetic stiffness with diabetic glucose levels (DsDg).

**Supplemental Fig. 2 Representative Trackmate videos.** Trackmate cell tracking was applied to the 48-hour time-lapse acquired with 10-minute intervals, from SPY-595 stained nuclei. Phase images were taken by a Keyence BZ-X800 fluorescence microscope with a 10x objective and a BZ-X Filter TexasRed filter. The two substrate stiffnesses are normal stiffness (Ns) at 57±5 kPa and diabetic stiffness (Ds) at 90±6 kPa. The two glucose conditions are normal glucose levels (Ng) at 5.5 mM, and diabetic glucose levels (Dg) at 11.1 mM. A) Trackmate video of cell nuclei migrating on normal stiffness with normal glucose levels (NsNg). B) Trackmate video of cell nuclei migrating on normal stiffness with diabetic glucose levels (NsDg). C) Trackmate video of cell nuclei migrating on diabetic stiffness with normal glucose levels (DsNg). D) Trackmate video of cell nuclei migrating on diabetic stiffness with diabetic glucose levels (DsDg).
