## Supplementary material for "The interplay between stiffness and hyperglycemia on diabetic foot ulcer wound closure": cellrose.m

```
%% Modified Code by: Nourhan Albeltagy
%           Department of Bioengineering, Temple University, Philadelphia, PA 19122, USA
% Modified from (Wind Rose,MATLAB) 

function cellrose = cellrose(celldir, cellspeed)
% CELLROSE Generates a polar histogram (rose plot) of cell movement directions and speeds.
% 
% INPUTS:
%   celldir   - A vector of cell movement directions in degrees.
%   cellspeed - A vector of cell movement speeds (e.g., in um/min).
%
% OUTPUT:
%   cellrose  - A 2D histogram matrix representing the percentage of cells in each direction and speed bin.
%
% If no output argument is provided, the function plots a stacked polar histogram.

%%

% Convert the cell direction from degrees to radians (required for trigonometric functions)
celldir = deg2rad(celldir);

% Validate the inputs to ensure they are numeric vectors with valid values
validateattributes(celldir, {'single', 'double'}, ...
                            {'vector', 'real', 'nonnan', 'nonempty', 'finite'}, ...
                             '', 'celldir', 1);
validateattributes(cellspeed, {'single', 'double'}, ...
                              {'vector', 'real', 'nonnan', 'nonempty', 'finite'}, ...
                               '', 'cellspeed', 2);

% Define the bin edges for direction (in radians) and speed
% Directions are divided into 15 bins (24-degree intervals)
directedges = deg2rad([0, 0:24:336, 360]);

% Speeds are divided into 5 bins: 0-0.2, 0.2-0.4, 0.4-0.6, 0.6-0.8, and >0.8
speededges = [0:0.2:0.8 Inf];

% Compute a 2D histogram of cell directions and speeds
% Rows correspond to direction bins, and columns correspond to speed bins
CellRose = histcounts2(celldir, cellspeed, directedges, speededges);

% Combine the first and last direction bins to account for circular data
CellRose = [CellRose(1,:)+CellRose(end,:); CellRose(2:end-1,:)];

% Normalize the histogram to represent percentages of total cells
total_cells = sum(CellRose, 'all');  % Total number of cells
CellRose = (CellRose / total_cells) * 100;  % Convert to percentages

% Compute cumulative percentages for stacked plotting
CellRoseStack = cumsum(CellRose, 2);

% If no output argument is provided, plot the data
if nargout == 0
    % Define the bin edges for the polar plot (16 bins for cardinal directions)
    cardinals = deg2rad(linspace(0, 360, 16));

    % Plot stacked polar histograms for each speed bin
    % Each speed bin is represented by a different color
    polarhistogram('BinEdges', cardinals, 'BinCounts', CellRoseStack(:, 5), 'FaceColor', [1.0 0.7 0.0]);  % Speed > 0.7 um/min
    hold on;
    polarhistogram('BinEdges', cardinals, 'BinCounts', CellRoseStack(:, 4), 'FaceColor', [1.0 1.0 0.0]);  % Speed 0.5-0.6 um/min
    polarhistogram('BinEdges', cardinals, 'BinCounts', CellRoseStack(:, 3), 'FaceColor', [0.0 1.0 0.0]);  % Speed 0.3-0.4 um/min
    polarhistogram('BinEdges', cardinals, 'BinCounts', CellRoseStack(:, 2), 'FaceColor', [0.5 0.8 0.9]);  % Speed 0.1-0.2 um/min
    polarhistogram('BinEdges', cardinals, 'BinCounts', CellRoseStack(:, 1), 'FaceColor', [0.1 0.1 1.0]);  % Speed < 0.1 um/min

    % Set transparency for all histogram layers
    set(allchild(gca), 'FaceAlpha', 1);

    % Dynamically set the radial axis limit based on the maximum percentage
    max_percentage = max(CellRoseStack, [], 'all');  % Find the maximum percentage value
    rlim([0 15]);  % Set radial axis from 0% to 15%

    % Customize the plot appearance
    ax = gca;  % Get the current axes
    ax.RColor = 'k';  % Set radial axis gridline color to black
    ax.GridColor = 'k';  % Set gridline color to black
    ax.RAxisLocation = 90;  % Place radial axis labels at 90 degrees
    ax.RAxis.TickValues = 0:10:ceil(max_percentage / 10) * 10;  % Set radial axis tick values
    ax.RAxis.TickLabelFormat = '%g%%';  % Format radial axis labels as percentages
    ax.RAxis.FontSize = 20;  % Set font size for radial axis labels
    ax.LineWidth = 2;  % Set line width for radial axis
    ax.ThetaAxis.FontSize = 20;  % Set font size for angular axis labels

    % Add a legend to describe the speed bins
    legend('> 0.7 um/min', '0.5 - 0.6 um/min', '0.3 - 0.4 um/min', '0.1 - 0.2 um/min', '< 0.1 um/min');
end
end
```
