## Supplementary material for "The interplay between stiffness and hyperglycemia on diabetic foot ulcer wound closure": cellrose_script.m

```
%% Modified Code by: Nourhan Albeltagy
%           Department of Bioengineering, Temple University, Philadelphia, PA 19122, USA
% Modified from (Wind Rose,MATLAB) 

% cellrose function folder should be located in the same directory 

%%
clear

% Specify the filename and delimiter
name = 'Example';
filename_ang = [name, '_ang.txt'];  % Replace with your filename
filename_vel = [name, '_vel.txt'];  % Replace with your filename
delimiter = '\t';           % Replace with your delimiter (e.g., ',', ' ', '\t' for tab)

% Read the text file into a table
anglestable = readtable(filename_ang, 'Delimiter', delimiter, 'VariableNamingRule', 'preserve');
% Read the text file into a table
speedtable = readtable(filename_vel, 'Delimiter', delimiter, 'VariableNamingRule', 'preserve');

angles = anglestable{:,5};
speed = speedtable{:,3};

angles(isnan(angles)) = 0;
speed(isnan(speed)) = 0;

figure('Position', [100, 100, 800, 600])
cellrose(angles, speed)


%title(['Migration angle and velocity of ', num2str(length(angles)), ' cells'], FontSize= 20)
title('NsNg Migration angle and velocity', FontSize= 20)

saveas(gcf, [name,'_ang&v.jpg']);
```
