## Supplementary material for "The interplay between stiffness and hyperglycemia on diabetic foot ulcer wound closure": Tracks_code.m

```
%% Code by: Nourhan Albeltagy
%           Department of Bioengineering, Temple University, Philadelphia, PA 19122, USA

% This code is designed to take a xlsx folder in the format defined by (ibidi Chemotaxis and Migration Tool, https://ibidi.com/chemotaxis-analysis/171-chemotaxis-and-migration-tool.html)
% The output is a figure showing the migration tracks of individual cells and the center of mass
% The color code represents which quarter of the wound area the cell started from: red is the upper right corner, green is the upper left corner, blue is the lower right corner, and yellow is the lower left corner. A black X shows the center of mass (COM) 

%% Figure Formatting 

t = tiledlayout(1,1); % Create a tiled layout with one tile
limitnum = [-600 600]; % Set axis limits

% Configure the first set of axes
ax1 = axes(t);
axis([limitnum limitnum]) % Set axis limits
xticks(limitnum(1):200:limitnum(2)) % Set x-axis tick intervals
yticks(limitnum(1):200:limitnum(2)) % Set y-axis tick intervals
ax = gca; % Get current axes
ax.FontSize = 18; % Set font size for the axes
xlabel('X trajectory (um)') % Label for x-axis
ylabel('Y trajectory (um)') % Label for y-axis
grid on % Enable grid

% Configure the second set of axes
ax2 = axes(t);
ax2.Color = 'none'; % Make the second axes transparent
ax2.XAxisLocation = 'origin'; % Set x-axis location to origin
ax2.YAxisLocation = 'origin'; % Set y-axis location to origin
ax2.YLim = limitnum; % Set y-axis limits
ax2.XLim = limitnum; % Set x-axis limits
xticks(limitnum(1):200:limitnum(2)) % Set x-axis tick intervals
yticks(limitnum(1):200:limitnum(2)) % Set y-axis tick intervals
ax2.TickLength = [0 0]; % Remove tick marks
set(ax2,'xticklabel',[]) % Remove x-axis tick labels
set(ax2,'yticklabel',[]) % Remove y-axis tick labels
box on % Enable box around the axes

% Set figure position and size
x0 = 50; % X-coordinate of the figure window
y0 = 50; % Y-coordinate of the figure window
width = 700; % Width of the figure window
height = 700; % Height of the figure window
set(gcf,'position',[x0,y0,width,height]) % Apply figure position and size

%% Load variables 

xlsxtable = importdata("example.xlsx"); % Import data from the Excel file
trcs = xlsxtable.data; % Extract data from the imported table

% Initialize variables for normalization and plotting
plotnorm = {}; % Cell array to store normalized trajectories
j = 1; % Index for plotnorm
x = 1; % Index for trcsnorm
r = randi([50 100],1,1000)*0.01; % Generate random values for color intensity
trcsnorm = []; % Array to store normalized trajectory data
normx = trcs(1,4); % Initial x-coordinate for normalization
normy = trcs(1,5); % Initial y-coordinate for normalization
meanx = (max(trcs(:,4)) + min(trcs(:,4))) / 2; % Mean x-coordinate
meany = (max(trcs(:,5)) + min(trcs(:,5))) / 2; % Mean y-coordinate

%% Graphing loop 

for i = 1:length(trcs) 
    if i == length(trcs) % Skip the last iteration
    elseif trcs(i+1,2) == trcs(i,2) % Check if the next point belongs to the same trajectory
        trcsnorm(x+1,1) = trcs(i+1,4) - normx; % Normalize x-coordinate
        trcsnorm(x+1,2) = trcs(i+1,5) - normy; % Normalize y-coordinate
        x = x + 1; % Increment index for trcsnorm
    else 
        % Process and plot the current trajectory
        trcsnorm(:,2) = trcsnorm(:,2) * -1; % Flip y-coordinates
        plotnorm(j) = {trcsnorm}; % Store normalized trajectory
        hold on % Retain current plot
        
        % Determine color based on position relative to mean coordinates
        if normy <= meany
            if normx >= meanx
                plot(plotnorm{j}(:,1),plotnorm{j}(:,2),'LineWidth',0.65, 'Color', [r(j) 0 0]) % Red
            else
                plot(plotnorm{j}(:,1),plotnorm{j}(:,2),'LineWidth',0.65, 'Color', [0 r(j) 0]) % Green
            end
        else
            if normx >= meanx
                plot(plotnorm{j}(:,1),plotnorm{j}(:,2),'LineWidth',0.65, 'Color', [0 0.4 r(j)]) % Blue
            else
                plot(plotnorm{j}(:,1),plotnorm{j}(:,2),'LineWidth',0.65, 'Color', [1 r(j) 0.2]) % Orange
            end
        end
        
        % Reset variables for the next trajectory
        j = j + 1; % Increment index for plotnorm
        trcsnorm = []; % Reset trcsnorm
        x = 1; % Reset index for trcsnorm
        normx = trcs(i+1,4); % Update normalization x-coordinate
        normy = trcs(i+1,5); % Update normalization y-coordinate
    end
end 

%% Center of mass calculations 

n = length(plotnorm); % Number of trajectories
yi = zeros(1,n); % Initialize y-coordinates of endpoints
xi = zeros(1,n); % Initialize x-coordinates of endpoints

for ii = 1:n
    xi(ii) = plotnorm{1,ii}(end,1); % Extract x-coordinate of the endpoint
    yi(ii) = plotnorm{1,ii}(end,2); % Extract y-coordinate of the endpoint
end

% Calculate center of mass
xn = (sum(xi) / n); % Mean x-coordinate
yn = (sum(yi) / n); % Mean y-coordinate

% Plot center of mass
plot(yn, xn, Marker=".", MarkerSize=30, Color='black') 

%% End of graphing 

hold off % Release the plot
title(append('Tracks of ', num2str(length(plotnorm)), ' cells over 24 hours'), 'FontSize', 24) % Add title
```
